# Effects of faecal inorganic contents in accurate measures of stress and nutrition hormone in large felines: implications for physiological assessments in free-ranging animals

**DOI:** 10.1101/2020.11.05.370635

**Authors:** Shiv Kumari Patel, Suvankar Biswas, Sitendu Goswami, Supriya Bhatt, Bivash Pandav, Samrat Mondol

## Abstract

Non-invasive stress and nutritional hormones and their interactions are increasingly being used to monitor psychological and nutritional physiology in free-ranging animals at different ecological scales. However, a number of extrinsic and intrinsic factors including hormone-inert dietary materials, inorganic matters etc. are known to affect accurate hormone measures. Here we addressed the impacts of inorganic matter (IOM) on corticosterone and T3 measures in wild tiger (n=193 from Terai Arc landscape, India) and captive lion (n=120 from Sakkarbaug Zoological Garden, Gujarat, India) faeces and evaluated possible corrective measures. The wild tiger samples contained highly variable IOM content (9-98%, mostly with >40% IOM) compared to captive Asiatic lion (17-57%, majority with <40% IOM). We observed significant negative correlation between IOM content and tiger GC (r=−0.48, p=0.000) and T3 (r=−0.60, p=0.000) measures but not in captive lions (r=−0.05, p=0.579). Two corrective measures viz. removing samples with ≥80% IOM and subsequently expressing concentrations as per gram of organic dry matter (instead of total dry matter) reduced IOM influence on tiger GC and T3 measures without affecting lion GC results. The corrective measures bring out alterations in the tiger T3 results but no changes in GC results. As faecal IOM content is associated with specific behaviours of many carnivore species, our results emphasize the need to reduce IOM-driven hormone data variation for ecologically relevant interpretations for species conservation.

## 1. Introduction

The ever-increasing impacts of humans on wildlife through habitat destruction and other environmental insults call for various reliable indices that can monitor temporal and spatial variations in physiological responses of free ranging animals. Faecal hormone analysis has become a powerful tool in this regard due to the variety of stress, nutrition and reproductive measures that can be used to evaluate the magnitude and direction of physiological responses of disturbances (Creel et al., 2002; Gobush et al., 2008; Keay et al., 2006; Wasser et al., 1997, 2005, 2011, 2017). Such non-invasive approach allow easy access of samples at spatial and temporal scales and provide a cumulative and feedback-free measure of circulating hormone over longer time period (Brown et al., 1996; Graham and Brown, 1996; Palme et al., 1996, 2005) when compared with blood (Sheriff et al., 2010). The interactive effects of these stress, nutrition and reproductive hormones also allow us to comprehensively relate wildlife health measures and adaptations to environmental changes in response to various disturbances (Ayres et al., 2012; Corkeron et al., 2017; Gobush et al., 2008; Hayward et al., 2011; Hunt et al., 2006; Mondol et al., 2020). For example, glucocorticoid (GC) hormone rapidly increases in response to wide range of short or long-term psychophysiological stressors (Dallman and Bhatnagar, 2010; Wingfield and Romero, 2010), whereas triiodothyronine (T3), the bioactive thyroid hormone acts more slowly, altering metabolism in response to acute or chronic nutritional stress (Eales et al., 1988; Flier et al., 2000). Similarly, monitoring reproductive hormones along with the psychological and nutritional stress are critical in understanding how various pressures take their toll on animal’s reproductive status (Foley et al., 2001; Vynne et al., 2014) When studied together, they provide complementary information on the animal coping-up mechanisms under different ecological or environmental conditions at different scales.

Despite such potential of the non-invasive endocrinology approaches in wildlife conservation, accurate measurements of faecal hormones is often challenging. A myriad of extrinsic and intrinsic factors are known to affect accurate measures of various faecal hormones including sampling and storage methods (Hunt and Wasser, 2003; Khan et al., 2002; Lafferty et al., 2019; Millspaugh et al., 2003b; Wilkening et al., 2016), species and sex-specific hormone metabolite compositions (Brown et al., 1994; Goymann et al., 2012; Young et al., 2004), and appropriate laboratory protocols for hormone extraction and assays (Davidian et al., 2015; Gholib et al., 2018; Millspaugh et al., 2003a; Nugraha et al., 2017; Pappano et al., 2010; Watson et al., 2013). The influences of these major factors in physiological assessments and their technical solutions have been the major focus of research over years (Millspaugh and Washburn, 2004; Palme et al., 2013, 2019; Sopinka et al., 2015; Touma and Palme, 2005). However, alterations introduced by dietary differences in free ranging animals that changes faecal composition and output have received less attention (Goymann et al., 2012; Von der Ohe and Servheen, 2002). To this end, undigested parts of the diet (e.g., dietary fiber) and other inorganic matters (e.g., urates, soil) that do not contribute hormone metabolite have been recognized to influence estimates of multiple steroid hormones in various species (Ganswindt et al., 2012; Wasser and Hunt, 2005). Studies suggest that these hormone-inert parts of faecal matter decreases hormone metabolite concentrations during extraction (Wasser and Hunt, 2005) and introduce errors in measuring stress (for example, see Ganswindt et al., 2012 for aardwolves; Goymann, 2005 for stone chats; Wasser and Hunt, 2005, Hayward et al., 2010 for owls etc.) and reproductive hormones (for example, see Goymann, 2005 for stone chats; Wasser et al., 1993 for baboons; Wasser and Hunt, 2005 for owls etc.). Such effects seem to be more pronounced in free-ranging animals, possibly due to large variations in species-specific food quantity and quality in the wild compared to captivity (Dierenfeld et al., 1994; Tilson et al., 2016). If unaccounted, these variations can result in inaccurate physiological interpretations resulting in misguided conservation efforts.

In this paper, we addressed key methodological issues to address the impacts of hormone-inert inorganic matters on measures of stress (corticosterone) and nutritional (T3) hormones in wild tigers in the Terai-Arc landscape, north India. The tiger (*Panthera tigris*) typifies global international conservation efforts across their range countries (Sanderson et al., 2010). Decades of intense conservation efforts have resulted in doubling their population in India (Jhala et al., 2015, 2020), and their future persistence will depend on managing these increasing population within the existing habitats (Gubbi et al., 2016, 2017). It is expected that the increasing tiger density will have distinct physiological impacts, particularly psychological stress (from intra-specific competition and anthropogenic disturbances), nutritional stress (wild prey depletion) and possibly reproductive effects. Tigers occupy a wide variety of habitats (Jhala et al., 2020) with varying dietary regimes (Basak et al., 2016; Harihar et al., 2011; Kumar et al., 2008) and their faeces is known to contain significant amount of soil and other inorganic matter (Khan, 2004; Schaller, 1967), thereby making this an informative system to study impacts of these materials on hormone measurements. While other studies have focused on stress measurements in wild tigers in India (Bhattacharjee et al., 2015; Malviya et al., 2018; Naidenko et al., 2019; Tyagi et al., 2019), they have not looked at the impacts of inorganic matters on stress or nutrition measures. In addition, we used another large feline carnivore species, the Asiatic lions (*Panthera leo persica*) from captivity where differences in dietary regimes are generally negligible and hence variations in inorganic matter content and its resulting influence on hormone measurements are expected to be low. Our main objectives were: (1) estimate the amount of inorganic matter present in faecal samples from wild tigers and captive lions and evaluate their variations, if any; (2) assess the impacts of such inorganic matter in measures of corticosterone (stress) and T3 (nutrition) hormones; and (3) evaluate how such errors affect the results and explore approaches to minimize them. We believe that our results have wider relevance for non-invasive endocrinology studies of wild animals, particularly carnivores with variable inorganic matter contents in their faeces.

## 2. Materials and Methods

### 2.1 Research permissions and ethical considerations

All required permissions for field surveys and biological sampling were provided by Forest Departments of Uttarakhand (Permit no: 90/5-6), Uttar Pradesh (Permit no: 1127/23-2-12(G) and 1891/23-2-12) and Bihar (Permit no: Wildlife-589), respectively. Due to non-invasive nature of work, no further ethical clearance was required for this study.

### 2.2 Study area

We sampled five major protected areas in the Indian part of Terai-Arc landscape (TAL) to collect wild tiger faeces. This linear landscape of TAL contains a total of 20800 km^2^ of potential tiger habitat, covering the states of Uttarakhand, Uttar Pradesh and Bihar (Qureshi et al., 2006). Situated at the foothills of the Himalayas, the habitat supports tropical moist deciduous forests dominated by Sal (*Shorea robusta*), tall Terai swamp grasslands and permanently moist reed swamps (Champion and Seth, 2005). This landscape is identified as &#8216;Global priority’ tiger conservation landscape (Sanderson et al., 2006) and retains about 22% of the India’s wild tiger population (Jhala et al., 2020). Our sampling was mostly concentrated inside five main tiger reserves in this landscape: Rajaji and Corbett Tiger Reserves in Uttarakhand (western TAL), Pilibhit and Dudhwa Tiger Reserves in Uttar Pradesh (central TAL) and Valmiki Tiger Reserve in Bihar (eastern TAL).

The captive Asiatic lion faecal sampling was conducted at the largest Asiatic lion conservation-breeding center at Sakkarbaug Zoological Garden (SZG), Gujarat. SZG hosts the largest captive population (N= 60) with the highest reported number of wild founders (Srivastav et al., 2018). We collected faeces from 35 individuals housed at the off-display conservation-breeding center of SZG. The same samples were used in an earlier study to evaluate the effects of enrichment interventions on behaviour and stress measures (see details in Goswami et al., 2020). During the enrichment intervention, the animals were randomly assigned to control (n=16) and test (n=19) groups to ascertain the stress measures during pre- and post-enrichment periods.

### 2.3 Faecal sample collection, species identification and hormone extraction

A team of experienced field personnel surveyed forest trails of aforementioned protected areas (see Bhatt et al., 2020) and collected fresh tiger faeces during winter seasons of 2016-2018. Faecal samples were stored in −20^0^C freezer till further processing. To identify tiger faeces we used DNA-based approaches described in Biswas et al. (2019). In brief, we swabbed each sample twice and digested the swabs overnight at 56^0^C with 30μl proteinase K and 300μl ATL buffer. Subsequently, Qiagen DNeasy tissue DNA kit protocol was followed to extract DNA. Negative controls were included to monitor any possible contamination. Post-extractions, we used tiger-specific mitochondrial DNA markers (Tig490F/R and Tig509F/R from Mukherjee et al., 2007) to ascertain tiger faeces, and used only confirmed fresh tiger faecal samples for downstream hormone and inorganic matter analyses. We used the following 193 tiger faecal samples in this study: Rajaji Tiger Reserve (RTR, n=57), Corbett Tiger Reserve (CTR, n=50), Dudhwa-Pilibhit region (DTR-PTR, n=52) and Valmiki Tiger Reserve (VTR, n=34). For lions, we used a total of 120 fresh faecal samples collected from the SZG conservation breeding center over a period of three months. We collected two fresh faecal samples per week from each individual during the enrichment experiment duration (covering both pre- and post-enrichment periods). The samples were stored in in −20^0^C freezer till further processing.

For hormone extraction from tiger (n=193) and lion (n=120), we followed a modified hormone extraction protocol described in Wasser et al. (2010). Each frozen sample was pulverized and dried up to 72 hours prior to hormone extraction to control for moisture (Wasser et al., 1993). Dried samples were then sieved through a 0.5 mm steel mesh strainers to remove prey remains/other hard parts and obtain faecal powder. The dried faecal powder was thoroughly mixed and subsequently hormone extraction was performed by pulse-vortexing 0.1 grams of powder in 15 ml of 70% ethanol, followed by centrifugation at 2200 rpm for 20 min (Mondol et al., 2020; Wasser et al., 2010). The hormone extracts were collected in 2 ml cryochill vials (1:15 dilution) and stored in −20°C freezer till further analyses.

### 2.4 Estimation of inorganic/organic content

To estimate inorganic/organic content in each faeces, we used a slightly modified protocol described in Ganswindt et al. (2012). We measured 0.1 grams (same weight as used for extraction) of dried faecal powder in a crucible and ashed the powder in a muffle furnace (#NSW-101, NSW, New-Delhi, India) at 550°C for 2 hours. Post-combustion, the residual inorganic matter (henceforth IOM) was weighed and amount of organic matter (OM) combusted in each sample was calculated using the formula: Organic matter = (0.1- weight of inorganic matter) grams.

### 2.5 Corticosterone and T3 assays

We used Corticosterone EIA (#K014, Arbor Assays, MI, USA) for measuring glucocorticoid (GC) metabolites in tiger and lion faecal extracts. Triiodothyronine (T3) EIA (#K056, Arbor Assays, MI, USA) was used for measuring T3 metabolites in faecal hormone extracts of wild tigers only. The lion samples were not assayed for T3 as in captivity the animals received uniform quantity of food based on their requirements and hence we did not expect any variations in T3 titers. Sample extracts were air-dried inside an incubator (#ATI-117, Obromax, Delhi, India) and re-suspended in assay buffer as per required dilutions. Each sample was assayed in duplicate using respective kit protocols and the optical density was measured at 450 nm using GMB-580 automatic microplate ELISA plate reader (#GMB-580, Genetix Biotech Asia, New Delhi, India). Hormone metabolite concentration is interpolated using four parametric logistic (4PL) regression function of GraphPad prism version 5 (GraphPad Software, California, USA). Cross-reactivities of respective antibodies are listed in Table 1.

**Table 1:** Details of the faecal hormone assays conducted for tiger and Asiatic lion in this study.

| Hormone | Assay method | Dilution | Slope ($R^2$) | Inter-assay CV | Intra-assay CV | Cross-reactivity |
| --- | --- | --- | --- | --- | --- | --- |
| Corticosterone (Lion pool) | EIA | 1:60 | 0.92 (0.99) | 9.0 | 9.8 | 100% with corticosterone, 12.30% with Desoxycorticosterone, 2.30% with Tetrahydrocorticosterone and <1% with Aldosterone, Cortisol, Progesterone, Dexamethasone, Corticosterone-21-Hemisuccinate, Cortisone and Estradiol |
| Corticosterone (Tiger pool) | EIA | 1:100 | 1.18 (0.99) | 9.0 | 3.6 |  |
| Triiodothyronine (Tiger Pool) | EIA | 1:15 | 0.98 (0.99) | 6.2 | 8.8 | 100% with T3, 0.88% with thyroxine and less than 0.1% with reverse T3 (3,3',5'-Triiodo-L-thyronine) |

### 2.6 Parallelism and Accuracy

We used parallelism and accuracy tests for laboratory validation of GC (both tiger and lions) and T3 (only tiger) hormone antibodies. We conducted parallelism tests for GC using dilutions of pooled tiger and lion faecal extracts from multiple random samples (n=20) to assess reliable quantification of respective hormones at different concentrations and to find optimal dilutions for final assays (at 50% binding). For T3 the same was performed with tiger pool samples. For both hormones, we generated species-wise sigmoid curves as relative dose vs. percent bound hormones for the pools and the standards, where parallel slopes indicate better antibody binding at different concentrations. Subsequently, we conducted accuracy test for both assays to determine any interference during antibody interactions. Independently we spiked both hormone standards with equal volumes of diluted faecal extract of known concentration (dilution level close to 50% bound from parallelism test) and assayed with standards. Results were plotted as regression lines using observed and expected concentrations from accuracy tests to show that faecal components were not interfering with assay accuracy at the tested dilution. Inter- and intra- assay coefficients of variation were determined using repeated measures of same-pooled extract.

### 2.7 Statistical analysis

Parallelism results were examined using an F ratio test for differences in slopes. Accuracy results were evaluated using analysis of linear regression. All analysis for parallelism and accuracy were performed in GraphPad prism version 5 (GraphPad Software, California, USA). Prior to further analysis, we assessed both GC (tiger and lion) and T3 (tiger) data for normality (both raw as well as log-transformed values) using diagnostic plots (Q-Q plots) and Shapiro-Wilk test. To assess the extent of IOM presence in faecal samples, we categorized them into five intervals of increasing percentage of IOM as 0-20%, 20-40%, 40-60%, 60-80%, and 80-100 %. We used arcsine- and square root-transformed IOM percentages as a variable and tested the strength and direction of association between IOM content and hormone concentrations (GC and T3, expressed as total dry matter) using Pearson’s correlation test. Earlier studies have described the effects of sample mass (biased GC measures from less faecal sample amount) (Hayward et al., 2010) and also the impacts of IOM in faecal GC measures (Ganswindt et al., 2012). To evaluate any potential impacts of IOM presence in faecal hormone measure we dropped samples with high IOM content (≥80% IOM) (to reduce sample mass effect). Further, we also calculated hormone concentrations as per unit of organic dry matter. We performed linear regressions to test for relationships between IOM content and hormone metabolite concentrations for all samples and with both types of corrected data sets (dropping samples with ≥80% IOM and hormone concentrations expressed as organic dry matter). Finally, for field collected tiger samples we compared the GC and T3 hormone levels among the sampled tiger reserves (RTR, CTR, DTR-PTR and VTR as mentioned above) using one-way ANOVA (separately for total dry matter and organic dry matter, respectively) along with post-hoc comparisons (Tukey’s HSD test) to detect any potential alteration in results before and after implementing corrective measures.

For Asiatic lions, we compared the post-enrichment GC data between the control and test groups as it showed significant differences between them (Goswami et al., 2020). We used independent t-test to evaluate any changes in results between two different ways of hormone concentration expressions (per units of total dry matter vs. total organic matter). We also calculated the effect sizes (ω^2^ for ANOVA and Hedges g for t-test) for each method. During all analyses, differences were considered significant at alpha level 0.05. Analyses were performed in GraphPad prism version 5, SPSS version 20 (IBM, 2011) and R v3.5.2 (R Core Team, 2018) using package “ggpubr” (Kassambara, 2020).

## 3. Results

### 3.1 Parallelism and accuracy

Parallelism and accuracy tests for tiger GC and T3 metabolites indicated reliable measures across different concentration ranges. Serial dilutions of faecal extracts paralleled the standard curves (Figure 1). There were no differences between slopes of standard and pooled extract curves for GC (F_(1,12)_ = 2.01, P = 0.182) as well as for T3 (F_(1,11)_ = 0.62, P = 0.45), and their curves were significantly different in their elevation for GC (F_(1,13)_ = 97.20, P <0.0001) and T3 (F_(1,12)_ = 19.77, P <0.0008). Accuracy tests produced slopes of 1.06 and 0.98 at working dilution of 1:100 and 1:15 for GC and T3 (Figure 1), respectively, suggesting that faecal extracts did not interfere with their metabolite measurement precisions. Intra-assay coefficient of variation (CV) was 3.6 and 8.84, whereas inter-assay CV was 9.0 and 6.18 for GC and T3, respectively (Table 1). Similarly, for Asiatic lions GC parallelism and accuracy produced reliable measures, where parallelism graphs showed no slope differences (F_(1,10)_ = 2.06, P = 0.182) and the curve elevations were significantly different (F_(1,11)_ = 66.25, P = <0.0001). Accuracy graphs showed a slope of 0.92 (at 1:60 dilution). Intra and inter assay CV was 9.8 and 8.99, respectively (Table 1).

**Figure 1:**
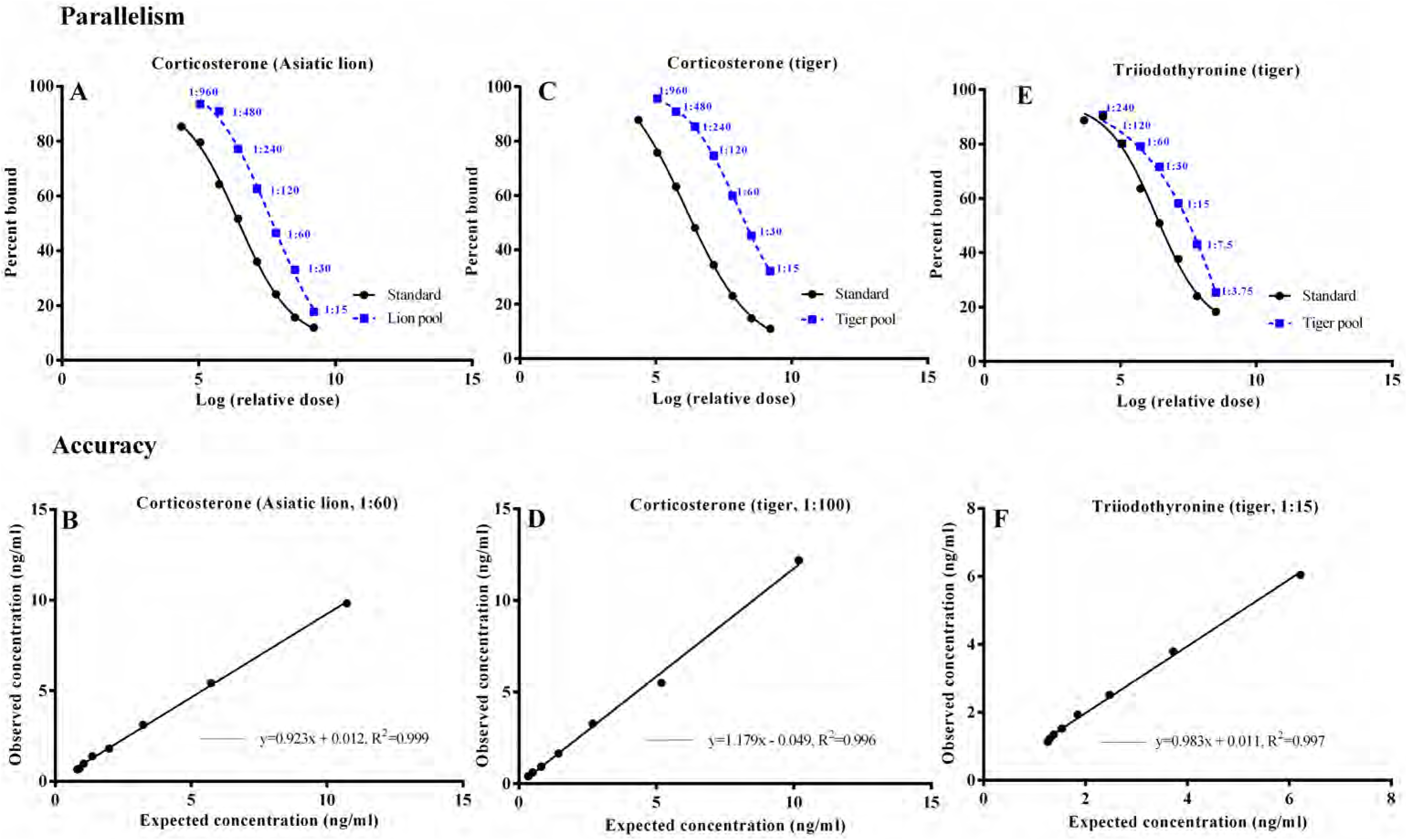
Parallelism and accuracy graphs for faecal corticosterone (both tiger and Asiatic lion) and T3 (tiger) EIA assays.

### 3.2 Impact of faecal inorganic content on hormone measure and alteration in results

We had a total of 193 and 187 samples data for GC and T3 analyses, respectively. Six tiger samples did not produce any data for T3 assays and were thus removed from analysis. For captive Asiatic lions, we had a total of 120 sample GC data from 35 different individuals. In terms of IOM content, the field-collected tiger samples showed higher variation (n=193, 9-98%) than the captive lion samples (n=120, 17-57%) (Figure 2). Majority of the tiger samples (n=178) had >40% IOM content, whereas IOM content remained <40% for majority of the lion samples (n=103). We observed a significant negative correlation between IOM content and GC (n=193, r=−0.48, p=0.000) and T3 (n=187, r=−0.60, p=0.000) metabolite concentrations from tiger samples, respectively whereas the lion samples with less variation in IOM did not show any such pattern (n=120, r=−0.05, p=0.579). Regression analysis with tiger samples showed strong influence of IOM content on GC measures when expressed as per gram of total dry matter (n=193, R^2^= 0.23, p= <0.0001) (Figure 3A). However, this influence reduces when samples with high IOM contents (≥80%) are removed from the analysis (n=139, R^2^= 0.06, p= 0.003) (Figure 3B), and reduces further when subsequently the data is expressed as organic dry matter (n=139, R^2^= 0.04, p= 0.013) (Figure 3C). Similarly, the T3 measures are also influenced strongly by IOM content when expressed as total dry matter (n=187, R^2^= 0.36, p= <0.0001) (Figure 3D). Removing ≥80% IOM samples reduced the influence (n=138, R^2^= 0.10, p= 0.0002) (Figure 3E), and subsequent expression of T3 concentrations as organic dry matter eliminates the influence of IOM (n=138, R^2^= 2.560E-4, p= 0.852) (Figure 3F). On the other hand, the captive lion samples showed no such influence of IOM content on GC measures when expressed as per gram total dry matter (n=120, R^2^= 0.003, p= 0.580) (Figure 5A) as well as per gram organic dry matter (n=120, R^2^= 0.017, p= 0.155) (Figure 5B). There were much less variation in IOM content and no samples were found with very high (≥80%) IOM proportion. Overall, these results suggest that field-collected samples have highly variable IOM contents that influence measures of faecal hormones, but these effects can be reduced by combining removal of samples with high IOM content and calculating hormone concentrations as per gram of organic dry matter.

**Figure 2:**
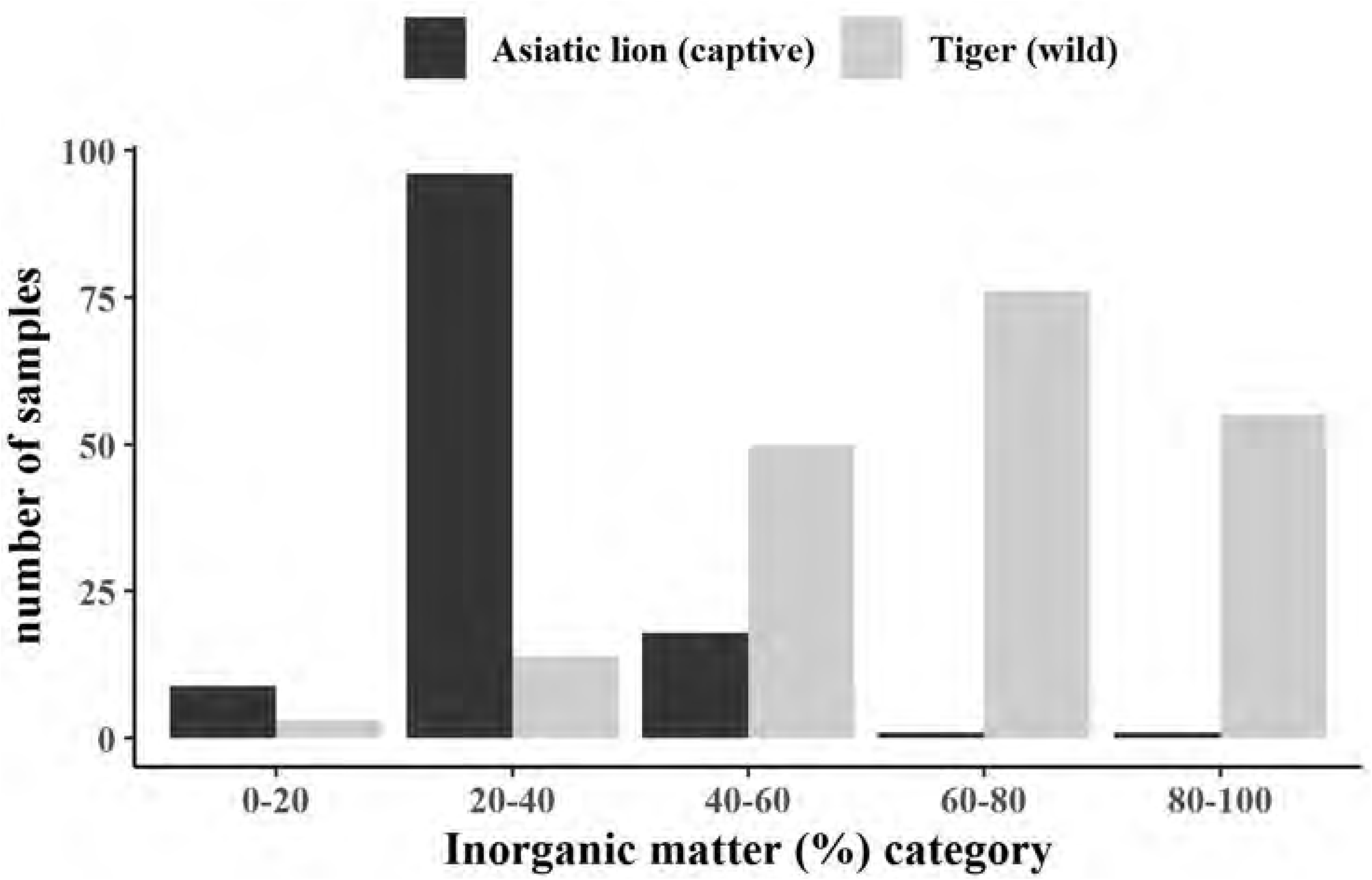
Comparison of Inorganic matter (IOM) content categorized in faecal samples of field-collected wild tigers and captive Asiatic lions. The samples were categorized into five groups.

**Figure 3:**
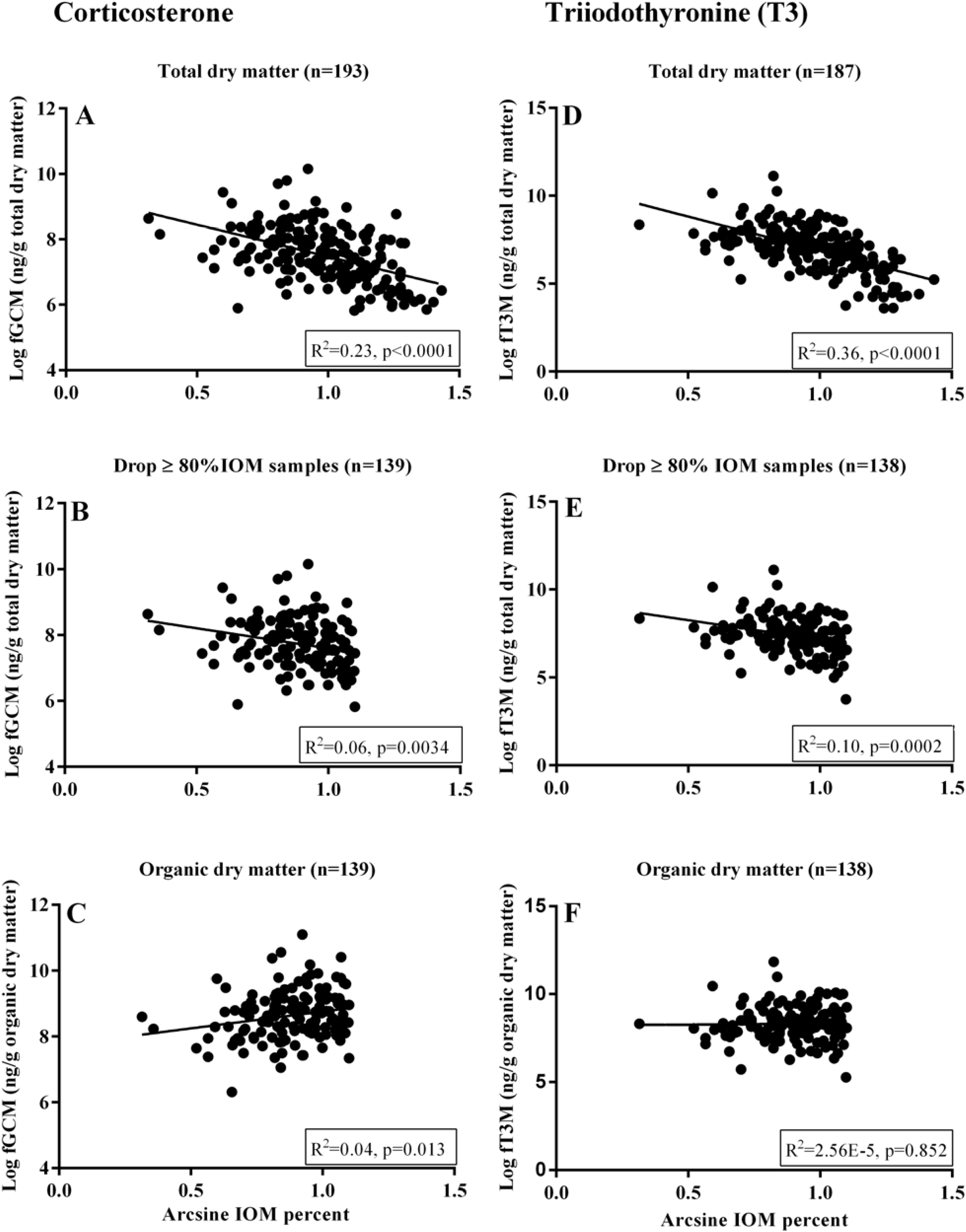
Relationship between arcsine-square root-transformed percentages of faecal IOM and log-transformed GC (fGCM) and T3 (fT3M) metabolite concentrations. Figure A and D represents results when hormone concentrations are expressed as per gram of total dry matter. Figure B and E shows results after dropping samples with high IOM contents (>80%). Figure C and F shows results when subsequently hormone concentrations are expressed as per gram of organic dry matter.

**Figure 4:**
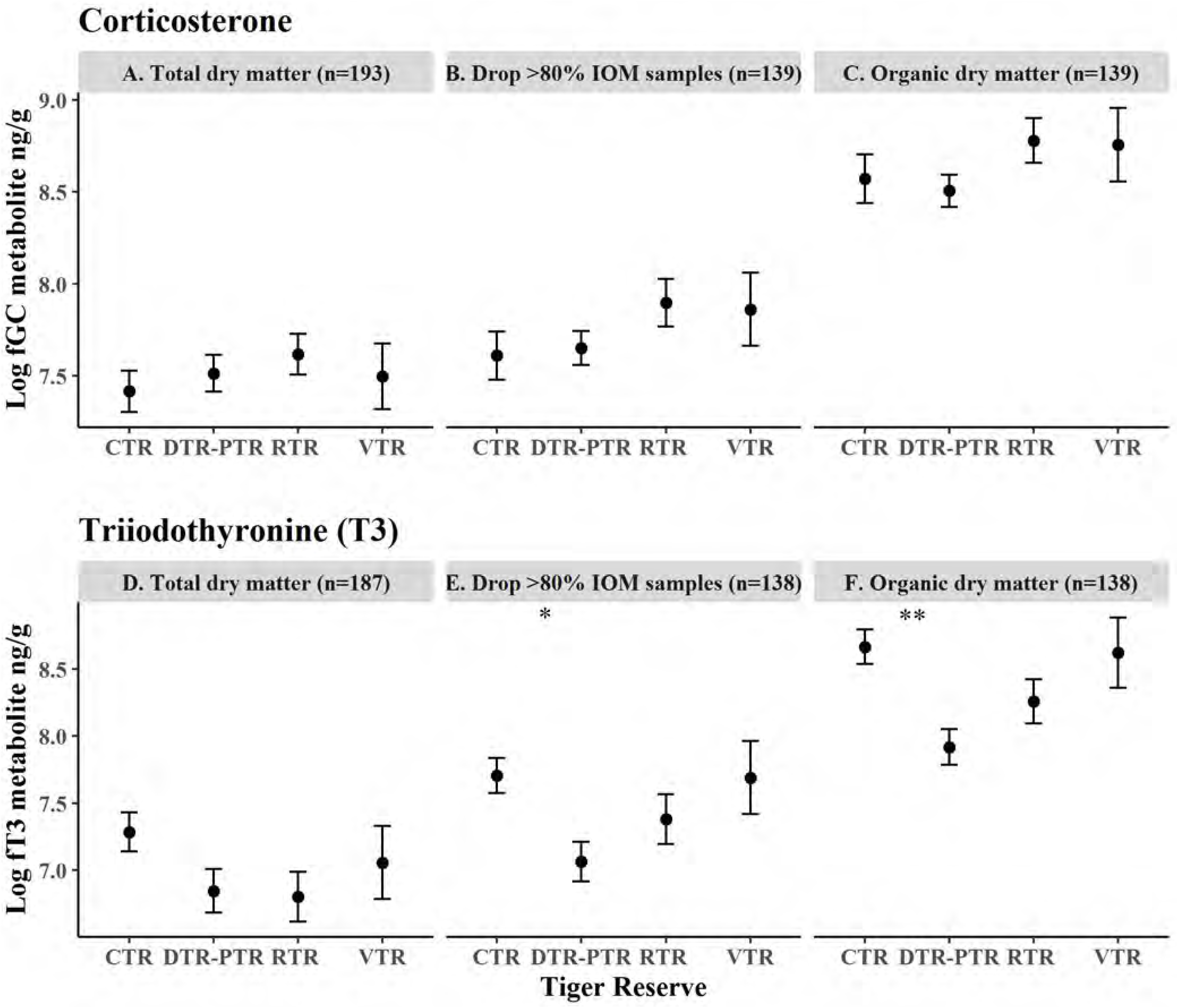
Comparison of tiger faecal corticosterone and T3 hormone concentrations (mean values) across four tiger reserves in Terai-Arc landscape (TAL). The top and bottom panel represents corticosterone and T3 data, respectively. For both hormones mean values (ng/g; log-transformed) are plotted. The left graphs (A and D) show the hormone concentrations expressed as per gram of total dry matter, the middle graphs (B and E) indicate patterns after dropping samples with high IOM contents (>80%) and the right graphs (C and F) represents patterns when hormone concentrations are expressed as per gram of organic dry matter. Any significant differences are marked with asterisks (*P< 0.05; **P<0.01).

**Figure 5:**
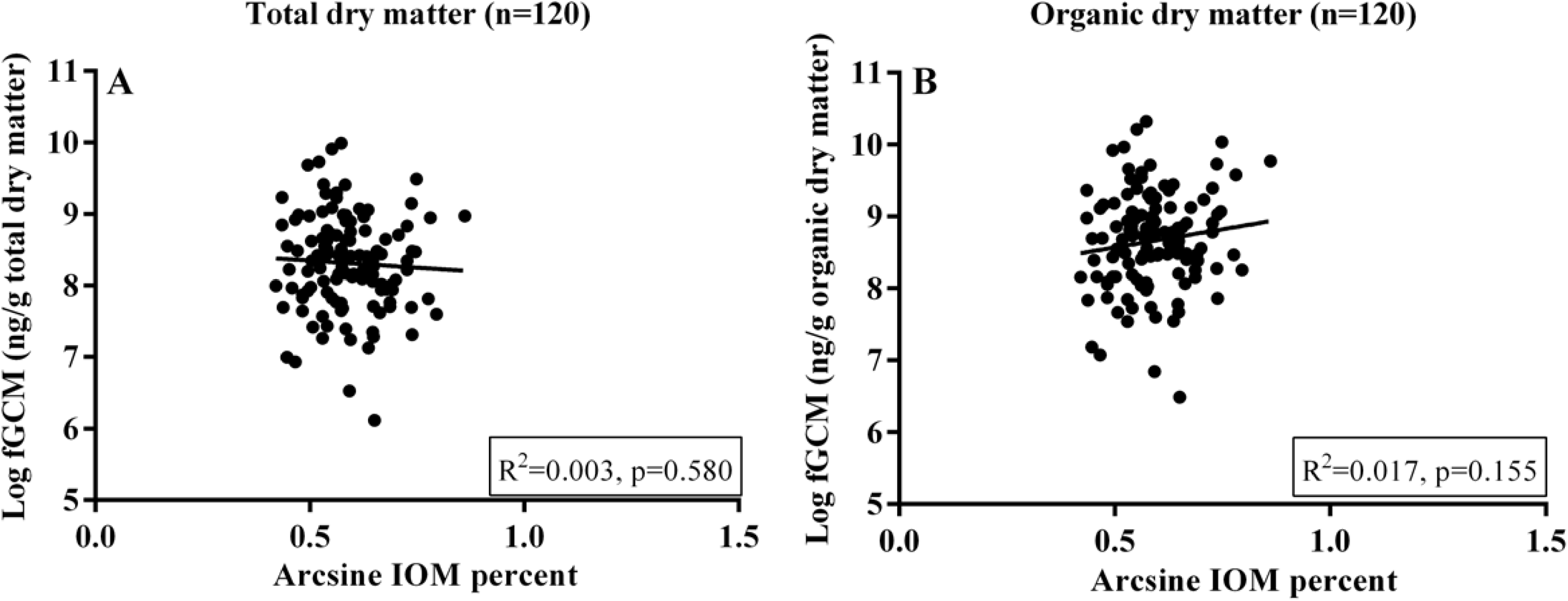
Relationship between inorganic matter (IOM) content and faecal corticosterone concentrations in captive Asiatic lions. The IOM values have been presented as arcsine-square root-transformed percentages and faecal hormone values have been log-transformed. The left graph (A) presents the association when hormone concentrations are expressed as per gram of total dry matter, whereas the right graph (B) indicates the association when hormone concentrations are expressed as per gram of organic dry matter.

One-way ANOVA analyses for field collected samples show no significant differences in mean GC levels across the sampled tiger reserves when concentrations were expressed as total dry matter (F (3,189)= 0.52, p=0.671, ω^2^ = −0.008), as well as when we removed samples with ≥80% IOM content (F (3,135)=1.26, p=0.290, ω^2^=0.007) and expressed the GC measures as organic dry matter (F (3,135)=1.18, p=0.322, ω^2^=0.004) (Figure 4A, B & C). However, we found significant changes in T3 results among three different datasets. We found no significant differences in mean T3 levels in sampled tiger reserves when concentrations were expressed as total dry matter (F (3,183)=1.56, p=0.200, ω^2^=0.009), but the values were significantly different when samples with ≥80% IOM content were removed from analysis ((F (3,134)=3.24, p=0.024, ω^2^=0.046). The significance increased further when T3 concentrations were expressed as per gram of organic dry matter (F (3,134) = 4.99, p=0.003, ω^2^ = 0.08) (Figure 4D, E & F). Subsequent Tukey’s HSD test for each of these two datasets show that CTR population has significantly higher T3 levels compared to DTR-PTR population (p=0.032, p=0.004, respectively) that has lowest T3 levels. Independent t-test results comparing GC levels between the post-enrichment control and test groups of captive lions showed non-significant changes in the concentrations (total dry matter (t(80)=9.77, p=0.00, g=2.21) and organic dry matter (t(80)=9.51, p=0.00, g=2.14) (Figure 6A & B), respectively. These analyses indicate that two different corrective measures for field collected samples bring out alterations in the tiger T3 results but no changes in GC results from four different protected areas. Single corrective measure for samples from captive Asiatic lions also showed no changes in the GC results.

**Figure 6:**
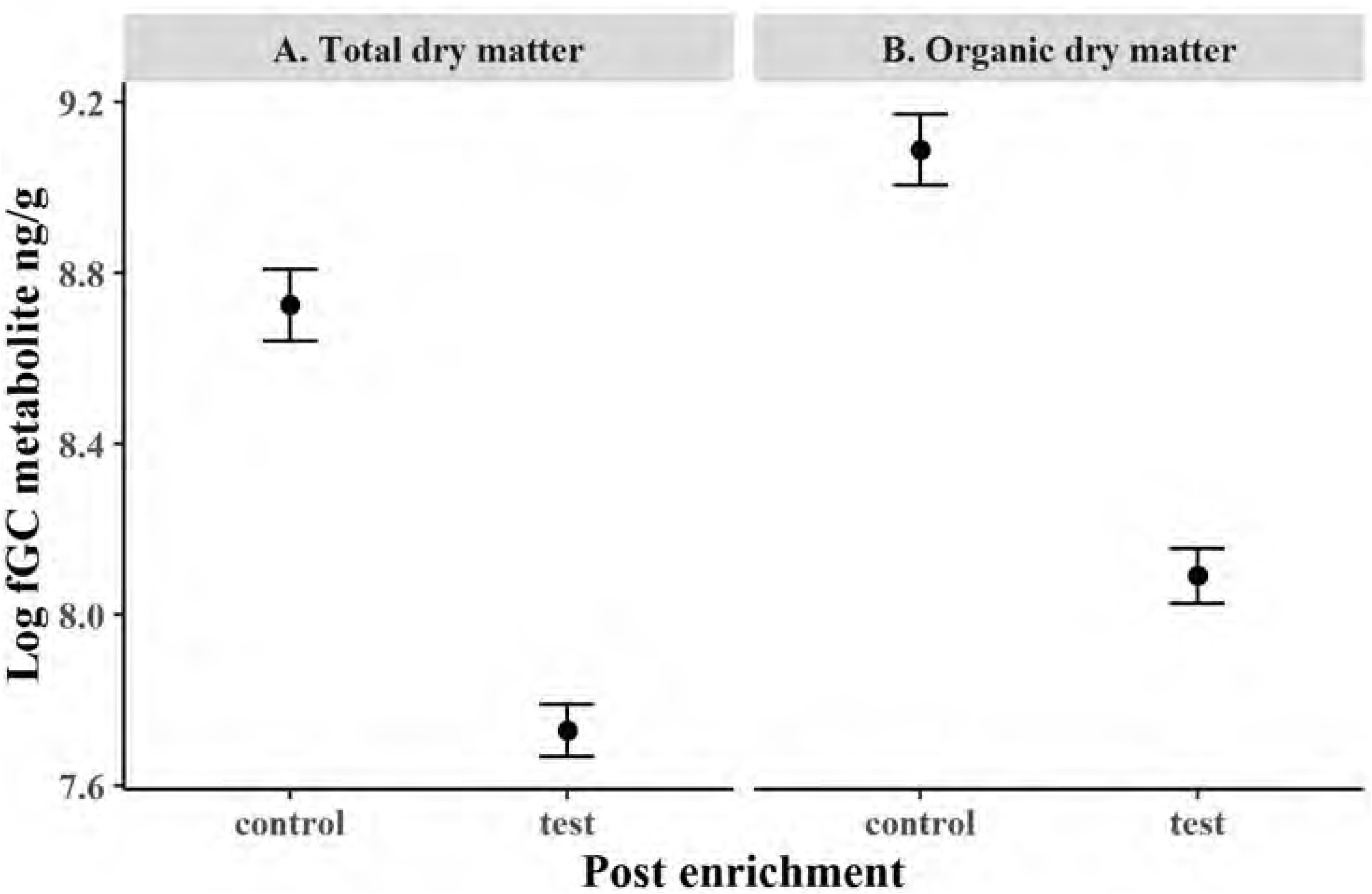
Comparison of captive Asiatic lion faecal corticosterone between control and test groups of animals after enclosure enrichment interventions, where mean corticosterone values (ng/g; log-transformed) are plotted. The left graph (A) shows the mean hormone concentration expressed as per gram of total dry matter, whereas the right graph (B) represent pattern when hormone concentration is expressed as per gram of organic dry matter.

## 4. Discussion

In this study we aimed to compare the impacts of inorganic matter (IOM) in measurement of faecal hormones from field-collected (tiger) and captive (Asiatic lion) animals. We found highly variable amount of IOM (9-98%) in the wild tiger faecal samples (n=193) where majority of the samples showed high (40 to ≥80%) IOM contents (Figure 2). However, the fecal samples of captive Asiatic lions on a standard diet (n=120) contained lower levels of IOM (20-40%) and lower variability among samples.. Although not quantified in this manner earlier, this pattern of IOM presence is not surprising for tigers. Khan (2004) reported presence of up to ~50% soil in wild tiger scats from the Sunderban landscape, Bangladesh during dry season with a peak in the winters and suggested possible seasonal soil ingestion by tigers. Schaller (1967) has also reported incidences of soil ingestion during winters from wild tigers in Kanha Tiger Reserve. However, such behaviours cannot always be attributed to specific seasons as free-ranging animals often naturally or deliberately ingest soil with food for mineral supplementation, alleviating gastrointestinal disorder or to counteract effects of high endoparasite load (Beyer et al., 1994; Knezevich et al., 1998; Krishnamani and Mahaney, 2000). Our sampling of tiger faeces during winters might have influenced the presence of IOM and their variations in the field-collected samples, but it is the best time for faecal sampling due to less leaf litter, availability of fresh samples and better environmental conditions. To the best of our knowledge, this is the first study on any endangered wild large carnivore species where such inter-sample variation in IOM is quantified. Earlier, Ganswindt et al. (2012) didn’t find significant inter-individual variation in IOM in Aardwolf scats, possibly due to low sample size (n=2) and similar foraging conditions in the study area. The possible presence of soil in tiger or other large carnivore faeces during sampling season and high variability of IOM makes it important to quantify its content before any downstream processing for hormone assessment.

Results of stress (GC) and nutrition (T3) metabolite measurements from the field-collected tiger faeces (with varied proportions of IOM) indicate a negative impact of IOM on both hormone concentrations (Figure 3A & D). However, the captive lion samples did not show any such pattern indicating that such effect in field collected samples is possibly due to dilution of hormone metabolites by IOM in faecal samples. This phenomenon has been earlier described in birds (Hayward et al., 2010) and mammalian species (Ganswindt et al., 2012). Taken together, we can infer that the hormone metabolites are mostly contributed by the organic part of the faeces and the use of best quality samples (in terms of organic matter content along with other parameters such as sample freshness) is critical for physiological assessments. Based on our results, we suggest two possible corrective measures to reduce the effects of IOM during hormone measures. First, is to explore possibilities of physical removal of IOM from faeces (as suggested for urates in case of birds by Hayward et al., 2010). However, such physical separation of IOM (soil, sand etc.) from field-collected faeces is difficult for carnivores and thus it is better to exclude the samples with high IOM content from analyses. Our data from tiger samples with high IOM (≥80%) content (and consequently with very low hormone contributing organic mass) show more variations in hormone concentration and removing them from analyses significantly reduces IOM influence on hormone measures (Figure 3B & E). Based on this, we suggest that samples with 80% or more IOM should not be processed for hormone assessments. We suggest a working protocol where the samples are first quantified for the IOM contents, followed by removal of samples with high IOM content and the downstream processes (for example, parallelism, accuracy etc.) are standardized with selected set of good quality samples. Future studies should also plan a pilot phase to evaluate the variation in faecal IOM in their respective target study systems before implementing a large-scale research project. Second, as these field collected samples still contain high variation in IOM amount, presenting the hormone titer on total dry matter units (per gram of total dry matter) can potentially be erroneous due to the dilutions from IOM rather than actual biological measures. However, expressing the hormone concentrations as per gram of organic dry matter has reduced the effects significantly in our study (Figure 3C & F), and supports the earlier suggestions by Hayward et al. (2010) and Ganswindt et al. (2012). For captive Asiatic lions where most of the samples were found with relatively low IOM quantity and the effects of these corrective measures are much less when compared to the field collected tiger samples. Overall, using a combination of removing very low organic matter-containing samples (≥80% IOM) and expressing the hormone concentration as per unit organic dry matter can help to reduce the negative influences of IOM on GC and T3 measures. Further work is required to check the IOM effects on reproductive hormone (progesterone, testosterone etc.) measures.

However, more experimental work is needed to assess the utility of the low-quality samples (in terms of IOM presence) for species with low densities and large home ranges, as collection of good-quality samples can often be challenging for them.

Finally, we investigated if these corrective measures mentioned above bring out any change in results by comparing the original and corrected tiger hormone data generated from five protected reserves. The mean GC levels of these tiger reserves remained non-significant for both original data (n=193 samples) (Figure 4A) as well as after both corrective measures (Figure 4B-C, Table 2). However, the T3 results across the three situations showed significant differences. For example, in the first scenario CTR and RTR showed highest and lowest mean T3 levels, respectively, with none significantly different from the other two populations (n=187, Figure 4D, Table 2). After dropping the samples with high IOM contents, mean T3 levels of DTR-PTR group drops to the lowest and the difference with CTR becomes statistically significant (n=138). Further, when the data is represented as organic dry weight (n=138, Figure 4E, Table 2) the significance remains with higher associated effect size (0.04 to 0.08) (Figure 4E, Table 2). The GC measures in captive Asiatic lions showed no difference in the result patterns between the original and corrected data (n=82, Figure 6A-B, Table 2). Since none of the Asiatic lion samples exceeded 80% IOM content we used only one corrective measure and the differences in mean GC concentrations between the control and test animals (post-enrichment) remained unchanged, differing only in titer values (Figure 6B).

**Table 2:**
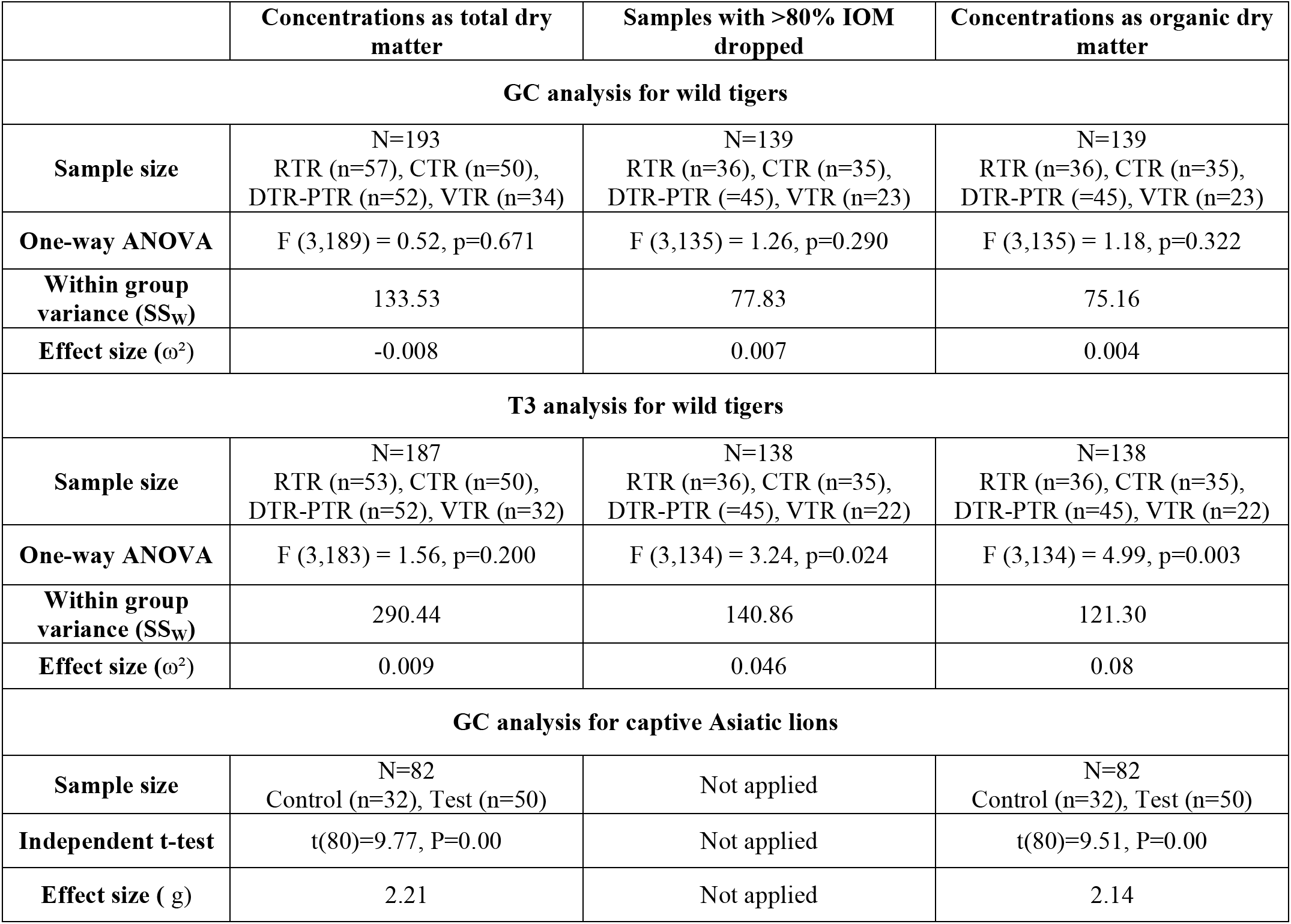
Comparison of hormone data analysis results for wild tigers and captive Asiatic lions. Wild tiger data was analyzed with One-way ANOVA and the captive Asiatic lion data was analyzed using independent t-test.

Our results show that certain common but under-emphasized factors such as IOM content have the potential to affect faecal hormone measures that in turn can change critical data interpretation and impact conservation decisions. Most of these physiological measures are closely associated with other ecological, behavioral and environmental factors at individual and population levels (Anestis 2010; Busch et al., 2009; Creel et al., 2013), but accurate interpretations should not be affected by measurement errors. We believe that the corrective measures discussed in this study reduce IOM-driven within-group hormone data variations and thus can help in making biologically relevant conservation decisions. Particularly for large, endangered carnivores such as tigers faecal sampling is the only non-invasive approach to assess species physiology in their natural habitats and thus it is essential to reduce the impacts of inorganic materials. These measures will help in accurate physiological assessments and lead us to ecologically relevant interpretations and recommendations, which are valued driving force for species conservation.

## 5. Acknowledgement

We acknowledge the Forest Departments of Uttarakhand, Uttar Pradesh and Bihar for providing necessary permits to carry out the research. Our thanks to the Forest Department officials and frontline staff members for their support and assistance during field surveys. We acknowledge help from Shrutarshi, Tista, Shrushti, Prajak, Harshvardhan, Rakesh, Sultan, Nimisha, Annu, Bura, Abbhi, Ranjhu, Ammi, Inam and Imam for their help during sampling and Chandra for assisting in laboratory work. We thank the Director, Dean, Research Coordinator and Nodal Officer of Wildlife Forensics and Conservation Genetics Cell of Wildlife Institute of India for their support.

This research was funded by Grant-in-Aid from Wildlife Institute of India. Samrat Mondol was supported by Department of Science and Technology INSPIRE Faculty Award (IFA12-LSBM-47).

## 6. Author contributions

**Shiv Kumari Patel:** Conceptualization, Data generation, Data curation, Formal analysis, Validation, Visualization, Writing-Original Draft, Writing-Review and Editing; **Suvankar Biswas:** Data generation, Data curation, Writing-Review and Editing; **Sitendu Goswami:** Data generation, Data curation, Writing-Review and Editing; **Supriya Bhatt:** Data generation, Data curation, Writing-Review and Editing; **Bivash Pandav**: Conceptualization, Resources, Writing-Review and Editing, Supervision, Project administration, Funding acquisition. **Samrat Mondol:** Conceptualization, Data Curation, Resources, Writing-Original draft, Writing-Review and Editing, Supervision, Project administration, Funding acquisition.

